# How does group differences in motion scrubbing affect false positives in functional connectivity studies?

**DOI:** 10.1101/2020.02.12.944454

**Authors:** Anders Eklund, Thomas E. Nichols, Soroosh Afyouni, Cameron Craddock

## Abstract

Analyzing resting state fMRI data is difficult due to a weak signal and several noise sources. Head motion is also a major problem and it is common to apply motion scrubbing, i.e. to remove time points where a subject has moved more than some pre-defined motion threshold. A problem arises if one cohort on average moves more than another, since the remaining temporal degrees of freedom are then different for the two groups. The effect of this is that the uncertainty of the functional connectivity estimates (e.g. Pearson correlations) are different for the two groups, but this is seldom modelled in resting state fMRI. We demonstrate that group differences in motion scrubbing can result in inflated false positives, depending on how the temporal auto correlation is modelled when performing the Fisher r-to-z transform.

## 1. Introduction

Motion scrubbing is commonly applied in resting state fMRI, to remove volumes where the head motion is greater than some threshold (Power et al., 2012). Motion scrubbing is performed independently for every subject, meaning that some subjects can have 90% of the fMRI volumes left after motion scrubbing, while other subjects may only have 50% of the volumes left. As an example, in the paper by Power et al. (2012) it is reported that 58% ± 20% and 26% ± 14% of the volumes were removed for two groups of children, but only 12% ± 6% of the volumes were removed for a group of adults. Siegel et al. (2014) reported 4% ± 5% scrubbing for typical adults and 16% ± 11% scrubbing for children. Similarly, Parkes et al. (2018) reported a significant difference in loss of temporal degrees of freedom between healthy controls and schizophrenics.

Group differences in motion scrubbing means that the average uncertainty of the functional connectivity estimates (e.g. Pearson correlations) can differ between the two groups, since the estimates are based on different number of measurements. Propagating subject uncertainty to the group analysis is, however, uncommon in resting state fMRI (an exception is the work by Fiecas et al. (2017)). This is in contrast to task fMRI, where the variance of each subject is commonly used in the group analysis (Woolrich et al., 2004; Chen et al., 2012), to for example downweight subjects with a higher variance. The problem of variable temporal degrees of freedom in resting state fMRI has been mentioned previously (Yan et al., 2013; Satterthwaite et al., 2013; Power et al., 2014; Pruim et al., 2015; Parkes et al., 2018), but to the best of our knowledge no study has investigated how it affects false positives in a group analysis. Here, we demonstrate that group differences in motion scrubbing can lead to inflated false positives rates for functional connectivity studies.

## 2. Data

We used the ABIDE preprocessed dataset (Di Martino et al., 2014; Craddock et al., 2013a)^1^, which contains preprocessed resting state fMRI data from 539 individuals diagnosed with autism spectrum disorder and 573 typical controls. The data were preprocessed using four different preprocessing pipelines (Connectome Computation System (CCS) (Xu et al., 2015), Configurable Pipeline for the Analysis of Connectomes (CPAC) (Craddock et al., 2013b), Data Processing Assistant for Resting-State fMRI (DPARSF) (Yan & Zang, 2010), NeuroImaging Analysis Kit (NIAK) (Bellec et al., 2011)), and are available for several preprocessing options (such as with and without bandpass filtering, and with and without global signal regression (GSR)^2^). Motion scrubbing has been applied for the NIAK pipeline, but not for the other pipelines, and we therefore used preprocessed data from CCS, CPAC and DPARSF. Mean timeseries were extracted for 7 regions of interest (ROI) atlases, and we focused our analyses on the Craddock 200 (CC200) atlas (Craddock et al., 2012), which consists of 200 ROIs. To lower the processing time, and to improve partial correlation estimates, we only performed the calculations for 50 of the 200 ROIs.

The ABIDE data were collected at 17 different sites, and all group analyses reported in this paper only used data from one site at a time. We focused on data collected at Michigan and New York, as these datasets contain the highest number of subjects (110 and 184, respectively). The downloaded data contains 82 subjects for Michigan and 171 subjects for New York, which is mainly explained by the fact that subjects with a mean frame-wise displacement larger than 0.2 are by default discarded by the ABIDE preprocessed download script. The Michigan datasets contain 296 volumes per subject, while the New York datasets contain 176 volumes per subject. All datasets were collected with a TR of 2 seconds.

## 3. Methods

### 3.1. Motion scrubbing

To investigate the effect of group differences in motion scrubbing on functional connectivity analyses, we used the same idea as in our previous work (Eklund et al., 2016, 2019); to perform many random group analyses with real fMRI data to empirically estimate false positives. Two random groups, of 20 subjects each, were first created by randomly shuffling all subjects and selecting the 40 first subjects. Preprocessed ABIDE data were then randomly motion scrubbed for every subject, where the proportion of scrubbing for each subject was randomly drawn from a normal distribution (at least 50 volumes were always saved). The mean proportion of scrubbing for the (fake) control group was set to 10%. For the (fake) diseased group, the mean proportion of scrubbing varied from 5% to 50%, in steps of 5% units. The standard deviation was set to 5% for the control group and 15% for the diseased group (Power et al., 2012). To reflect the fact that motion contaminated volumes often appear in clusters, rather than appearing randomly, a simulation script was used to create more realistic motion scrubbing, see Figure 1.

**Figure 1:**
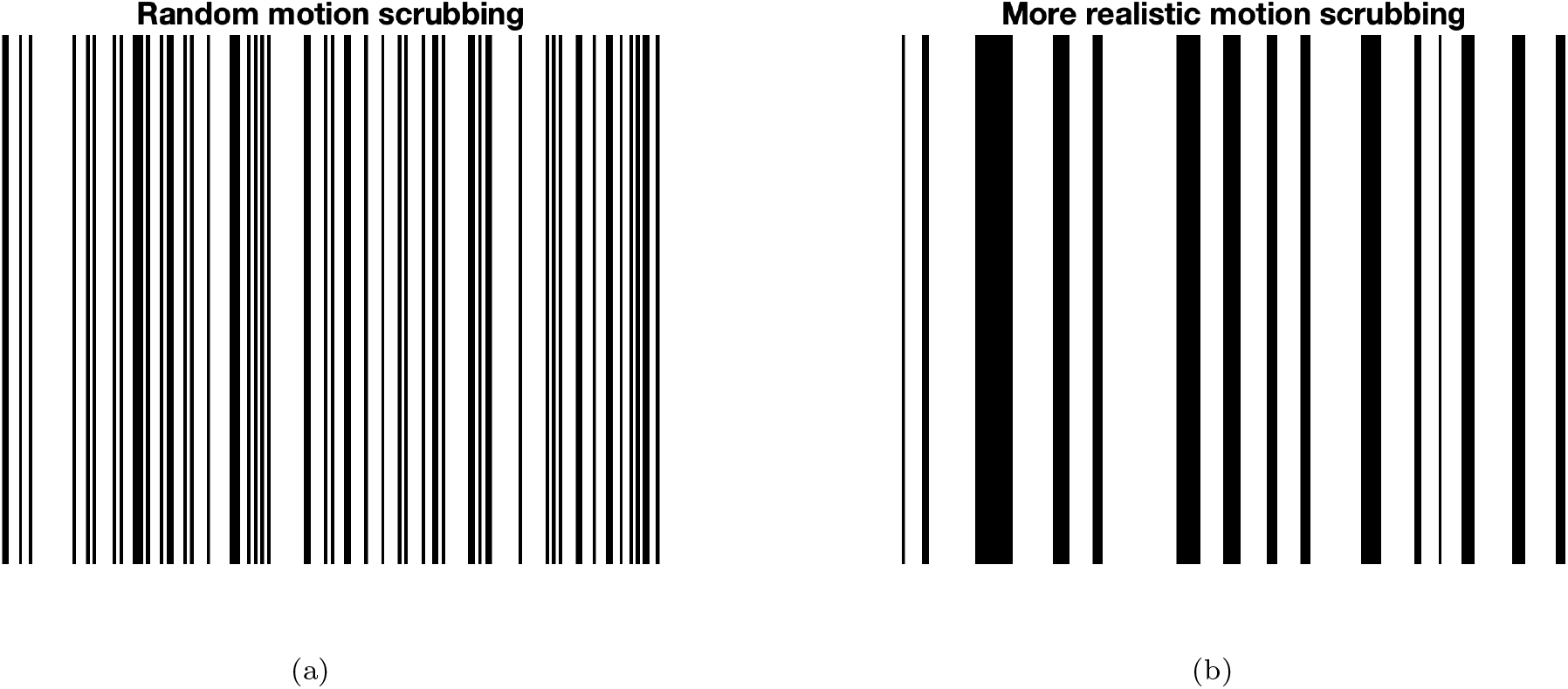
Comparing random scrubbing (left, obtained by the function randperm in Matlab) and more realistic scrubbing (right, obtained through a more advanced simulation) for a time series with 200 time points and 30% scrubbing (black time points represent time points to be removed).

### 3.2. ROI to ROI correlation estimation

Pairwise correlations (both full and partial) between all ROI time series were calculated using the motion scrubbed data (using the function nets netmats in FSLNets^3^ (Smith et al., 2011)). For partial correlation we used L2 norm regularization (Tikhonov), since L1 regularization is rather slow, and the default setting in FSLNets (0.1). In resting state fMRI the correlation values are normally Fisher r-to-z transformed before applying a t-test. Note that the Fisher transformation on its own will not produce z-scores, since it does not involve division with a standard error. The temporal auto correlation present in fMRI data complicates the variance estimation (Eklund et al., 2012; Arbabshirani et al., 2014; Afyouni et al., 2019), which can lead to inflated z-scores. FSLNets uses a global AR(1) model (the same for all ROI to ROI correlations) to model the temporal auto correlation. The AR parameter is estimated using a Monte Carlo simulation (and not through prewhitening (Bright et al., 2017; Honari et al., 2019)). Other softwares (such as REST (Song et al., 2011) and Conn (Whitfield-Gabrieli & Nieto-Castanon, 2012)) do not mention any model for the auto correlation. We therefore decided to perform the analyses with uncorrected (raw) and corrected (FSLNets global AR(1)) r-to-z transformation.

### 3.3. Group analysis

The group analysis consisted of applying a two sample t-test (using ttest2 in Matlab) to every element below the diagonal in the (r-to-z transformed) correlation matrix, to test if there is a difference in mean functional connectivity between the two randomly created groups (controls > diseased). Bonferroni correction was used to correct for multiple comparisons over correlation matrix elements, and should be conservative since the tests are not independent. A total of 96 different parameter combinations, listed in Table 1, were tested to understand how different parameters affect the results. The whole procedure was repeated 1000 times for every parameter combination, to empirically estimate the (familywise error corrected) degree of false positives (defined as the proportion of times a significant group difference was found, after correcting for multiple comparisons).

**Table 1:** Parameters tested for the different processing pipelines. One thousand random group analyses were performed for each parameter combination.

| Parameter | Values used |
| --- | --- |
| fMRI data | New York (171 subjects), Michigan (82 subjects) |
| Pipeline | CCS, CPAC, DPARSF |
| Bandpass filtering (0.01 - 0.1 Hz) | No, Yes |
| Global signal regression | No, Yes |
| Fisher r-to-z transform | Without autocorrelation correction (raw), With autocorrelation correction (AR(1)) |
| Analysis type | Full correlation, partial correlation (Tikhonov regularization) |

### 3.4. Reproducibility

As for our previous work (Eklund et al., 2015, 2016, 2019), our results are fully reproducible since we used open data and share the processing scripts on GitHub^4^. Other researchers can thereby reproduce our findings and extend the analyses to other settings (Eklund et al., 2017b; Poldrack et al., 2017).

## 4. Results

Figure 2 shows estimated FWE rates for different amounts of motion scrubbing per group and different preprocessing strategies, for the CCS pipeline and the New York data. Figures 3 and 4 show corresponding FWE rates for the CPAC and DPARSF processing pipelines. Figures 5 to 7 show corresponding results for the Michigan data. Clearly, using the r-to-z transform with auto correlation correction in FSLNets leads to inflated false positive rates for large group differences in motion scrubbing. Applying no correction for auto correlation leads to nominal results for all parameter combinations, and this is explained by the fact that the correlation (and the raw Fisher transformed correlation) are not test statistics, since they have not been divided by a standard error.

**Figure 2:**
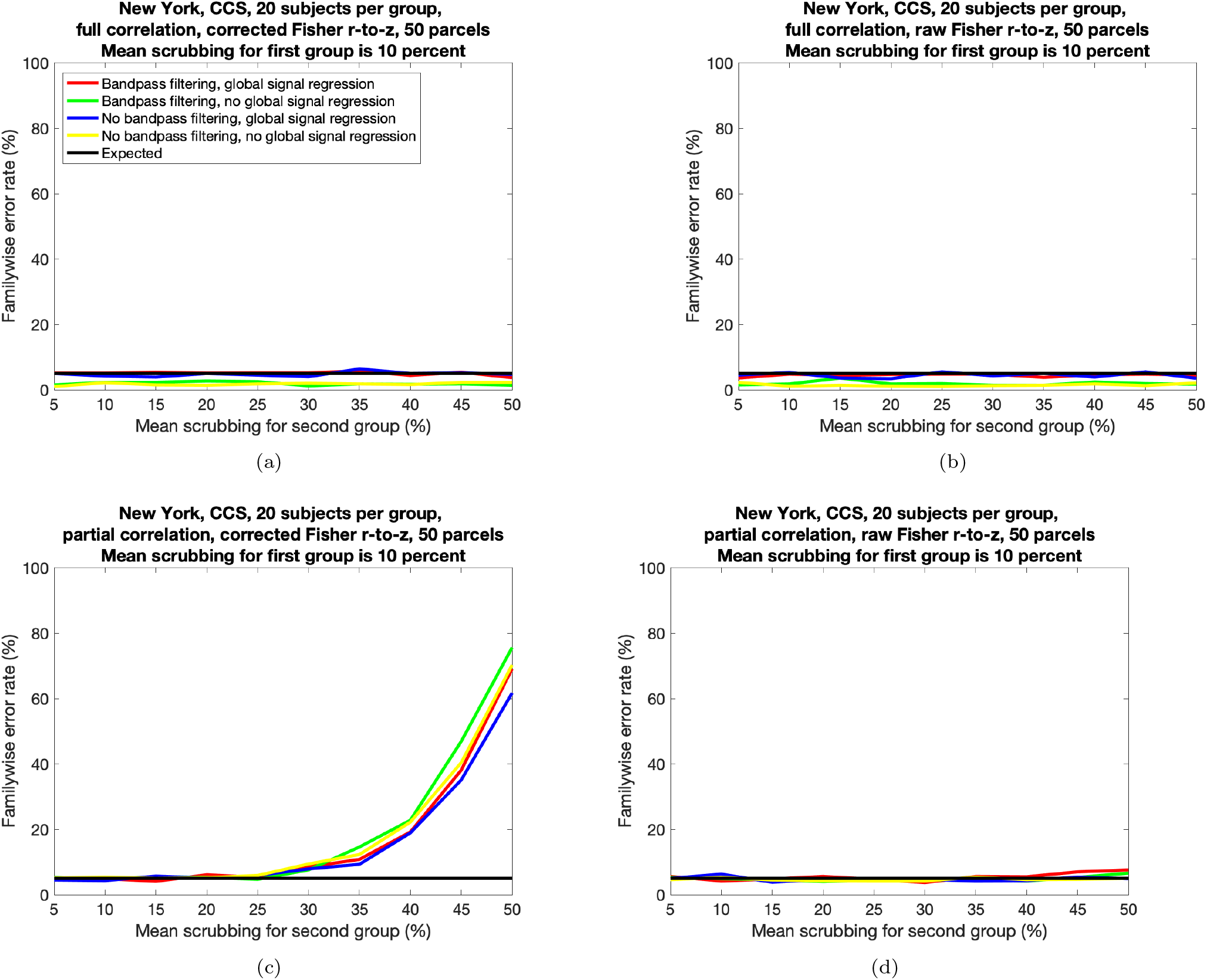
Estimated familywise error rates, for different proportions of motion scrubbing for two groups of randomly selected subjects, for the CCS processing pipeline. **Left:** Results with Fisher r-to-z transform with auto correlation correction **Right:** Results with Fisher r-to-z transform without auto correlation correction **Top:** Results for full correlation. **Bottom:** Results for partial correlation.

**Figure 3:**
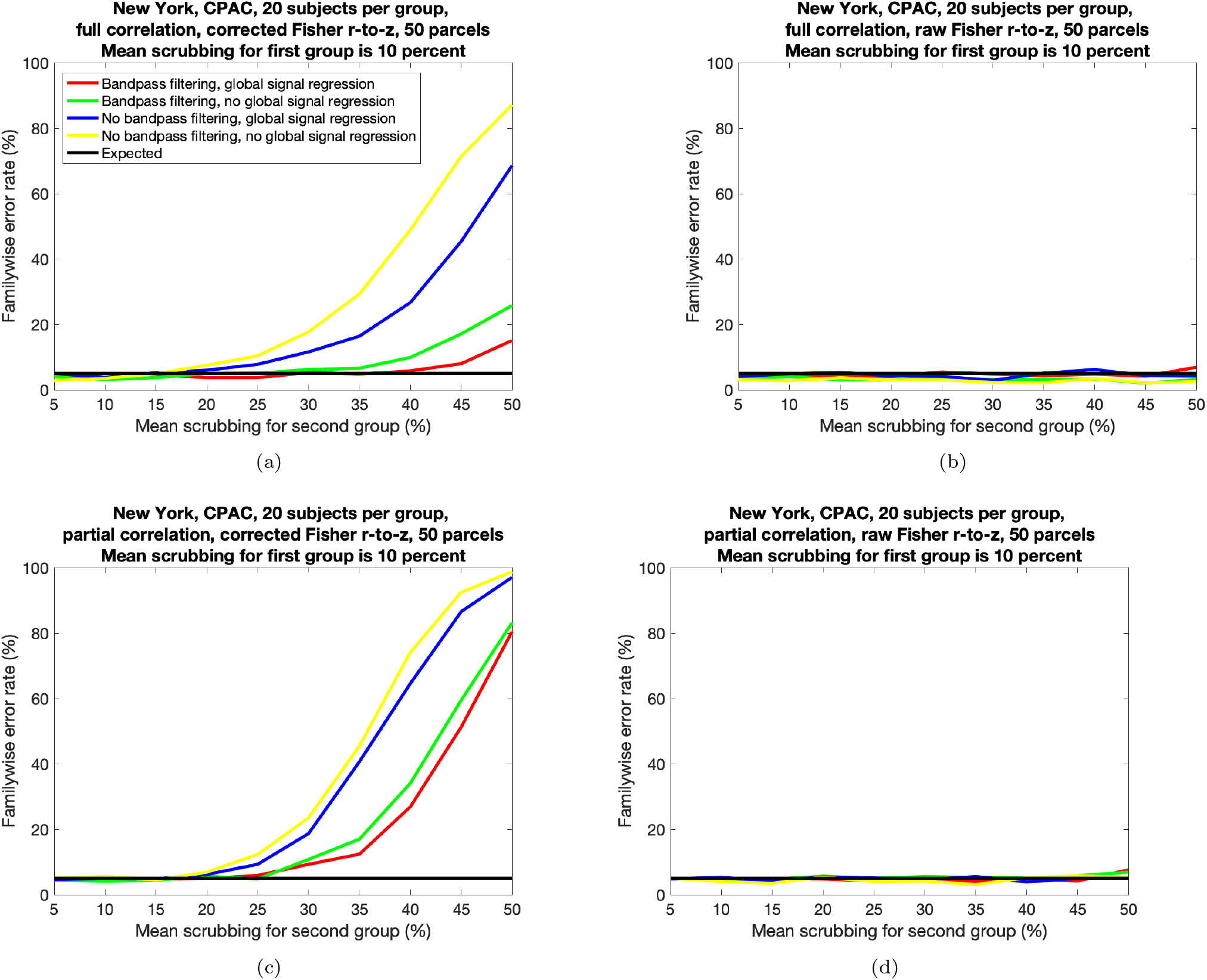
Estimated familywise error rates, for different proportions of motion scrubbing for two groups of randomly selected subjects, for the CPAC processing pipeline. **Left:** Results with Fisher r-to-z transform with auto correlation correction **Right:** Results with Fisher r-to-z transform without auto correlation correction **Top:** Results for full correlation. **Bottom:** Results for partial correlation.

**Figure 4:**
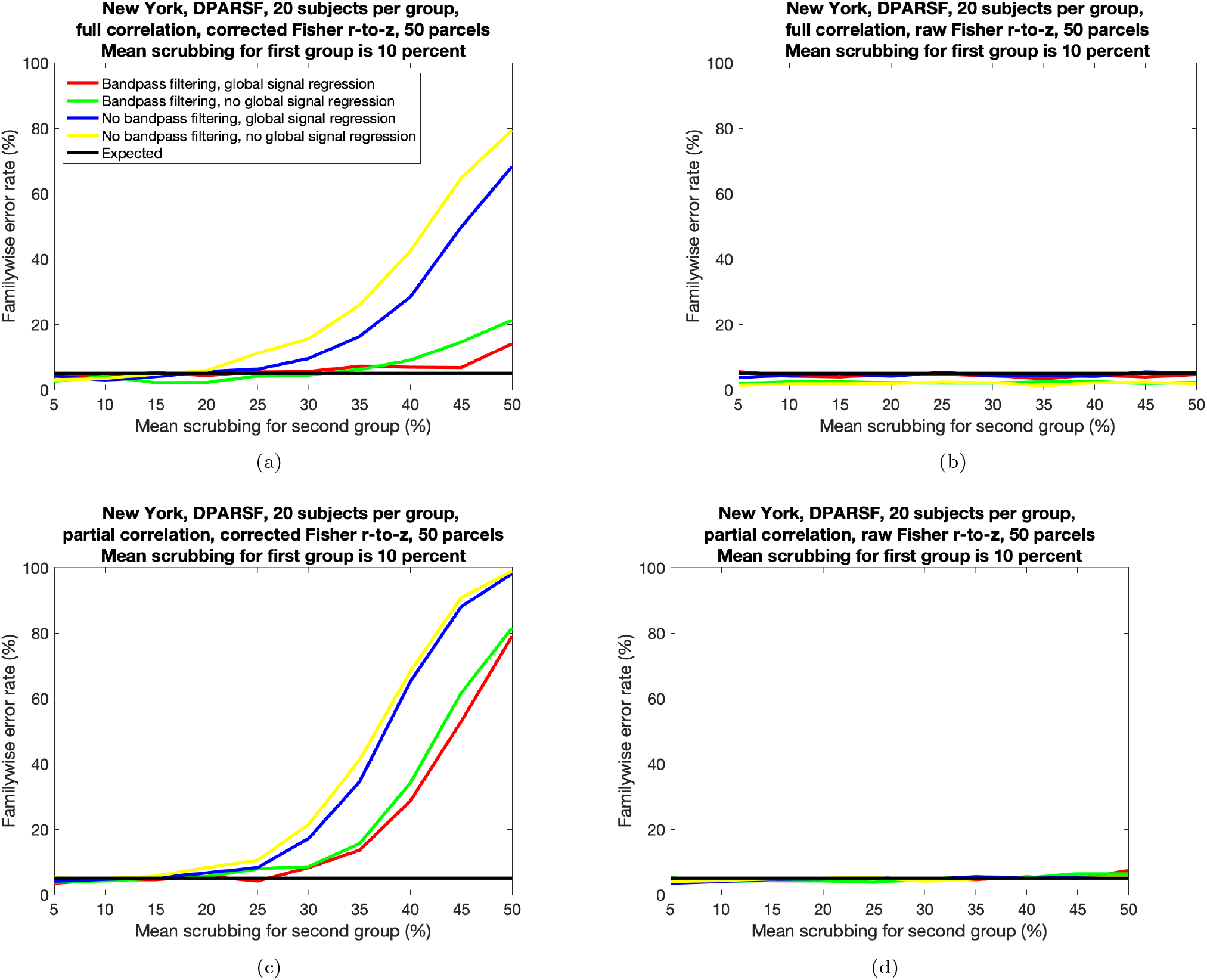
Estimated familywise error rates, for different proportions of motion scrubbing for two groups of randomly selected subjects, for the DPARSF processing pipeline. **Left:** Results with Fisher r-to-z transform with auto correlation correction **Right:** Results with Fisher r-to-z transform without auto correlation correction **Top:** Results for full correlation. **Bottom:** Results for partial correlation.

**Figure 5:**
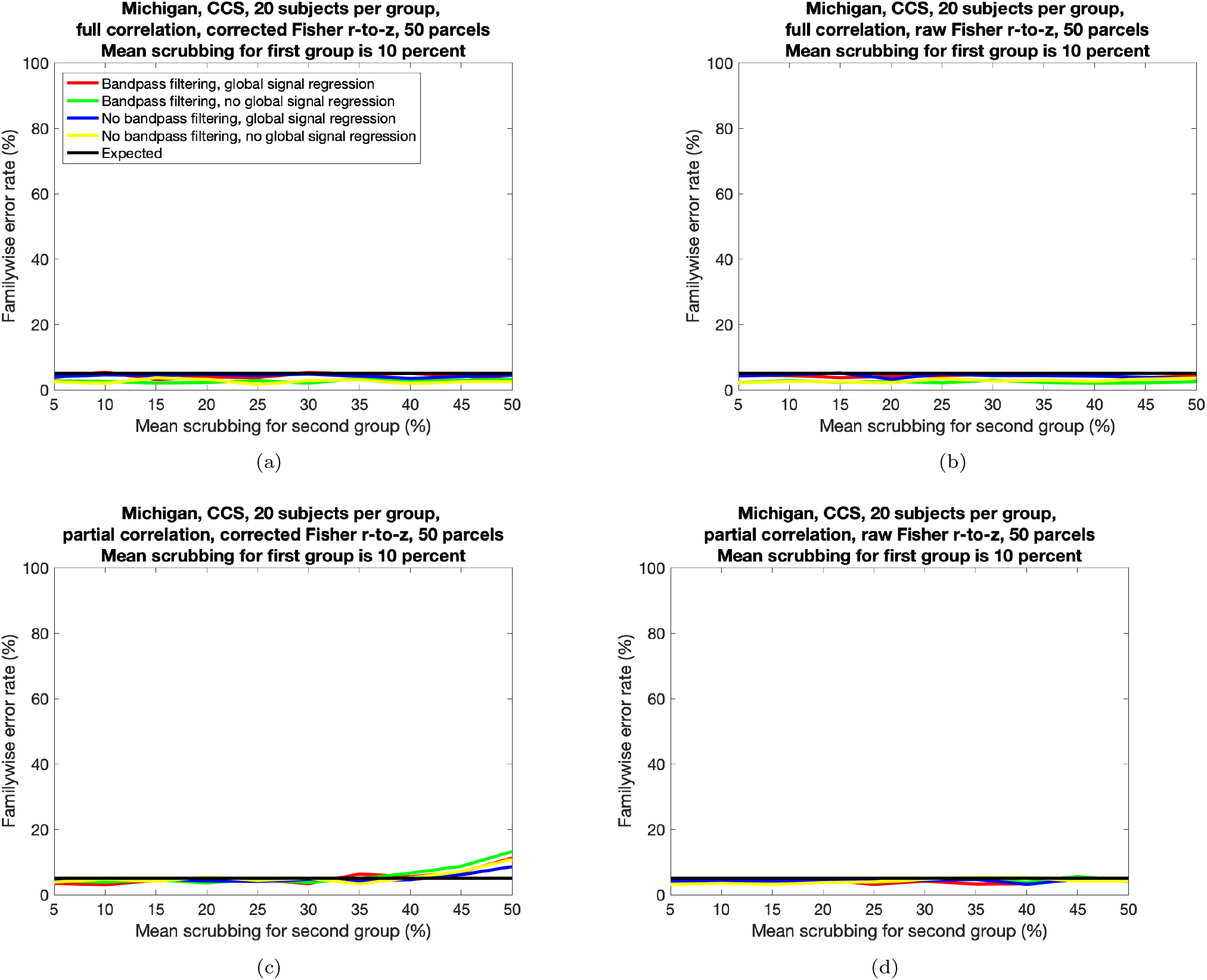
Estimated familywise error rates, for different proportions of motion scrubbing for two groups of randomly selected subjects, for the CCS processing pipeline. **Left:** Results with Fisher r-to-z transform with auto correlation correction **Right:** Results with Fisher r-to-z transform without auto correlation correction **Top:** Results for full correlation. **Bottom:** Results for partial correlation.

**Figure 6:**
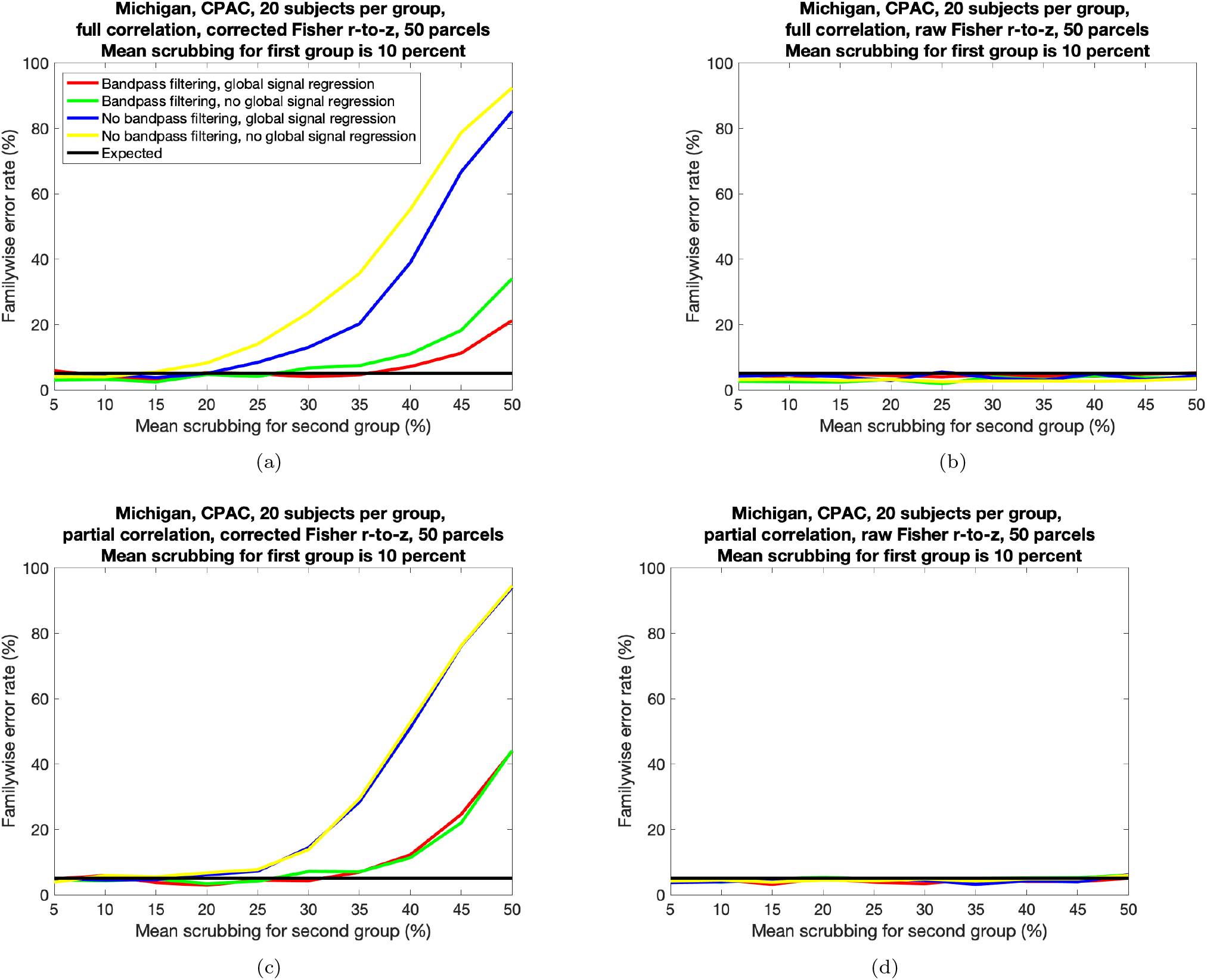
Estimated familywise error rates, for different proportions of motion scrubbing for two groups of randomly selected subjects, for the CPAC processing pipeline. **Left:** Results with Fisher r-to-z transform with auto correlation correction **Right:** Results with Fisher r-to-z transform without auto correlation correction **Top:** Results for full correlation. **Bottom:** Results for partial correlation.

**Figure 7:**
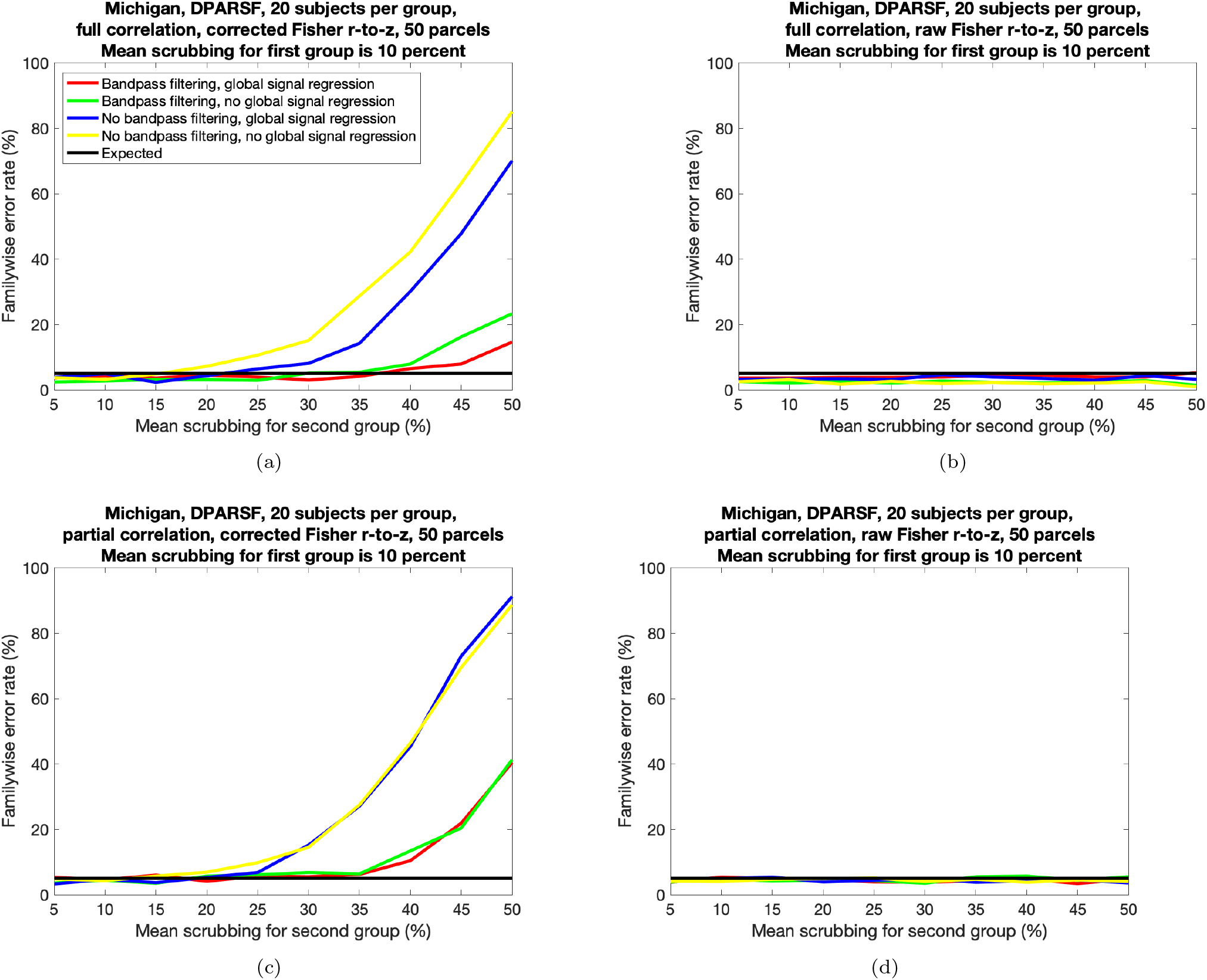
Estimated familywise error rates, for different proportions of motion scrubbing for two groups of randomly selected subjects, for the DPARSF processing pipeline. **Left:** Results with Fisher r-to-z transform with auto correlation correction **Right:** Results with Fisher r-to-z transform without auto correlation correction **Top:** Results for full correlation. **Bottom:** Results for partial correlation.

The FWE rate is in general lower when bandpass filtering is applied (mean 7.3%, compared to 12.5% without filtering). Global signal regression has a small effect on the mean FWE rate; 9.7% with GSR and 10.1% without GSR. The FWE rate is slightly higher for the New York data (mean 11.1%, compared to 8.8% for Michigan), which is expected since the Michigan data contains 296 time points per subject, compared to 176 time points per subject for the New York data. Furthermore, the FWE rate is in general higher for partial correlation (mean 12.3%, compared to 7.5% for full correlation). The mean FWE rate per pipeline is 5.7% (CCS), 12.5% (CPAC) and 11.5% (DPARSF). A possible explanation is that the CCS pipeline includes a despiking step, which will reduce ROI correlations due to spikes (as a spike will be present in most ROIs).

## 5. Discussion

We have presented results showing that group differences in motion scrubbing can lead to inflated false positive rates for functional connectivity studies, depending on how the r-to-z transform is performed. The problem of varying temporal degrees of freedom has been mentioned previously (Yan et al., 2013; Satterthwaite et al., 2013; Power et al., 2014; Pruim et al., 2015; Parkes et al., 2018), but we are not aware of any study that investigates how it affects false positives. Yendiki et al. (2014) investigated how group differences in motion affect diffusion MRI results, but did not use motion scrubbing or censoring. We have here focused on scrubbing (Power et al., 2012), but the results may also apply for spike regression (Satterthwaite et al., 2013) since adding one covariate per censored volume will also reduce the temporal degrees of freedom. It is not clear if scrubbing and spike regression would have the same effect on the autocorrelation structure of the time series. Instead of completely removing time points with too large motion, a heteroscedastic model can be used to automatically downweight affected time points (Eklund et al., 2017a). As in task fMRI, the uncertainty of each subject should also be propagated to the group analysis (Woolrich et al., 2004; Chen et al., 2012; Fiecas et al., 2017), to downweight subjects with a higher uncertainty.

### 5.1. False positives

Our results show that group differences in motion scrubbing only leads to inflated false positive rates if the z-score calculation in FSLNets accounts for the temporal auto correlation. As mentioned in the Methods section, the reason for this is that the correlation (and raw Fisher transformed correlation) are not test statistics, since they have not been divided by a standard error. The raw Fisher r-to-z transform, used in REST (Song et al., 2011) and Conn (Whitfield-Gabrieli & Nieto-Castanon, 2012), does not consider the number of time points in the time series, and will therefore produce the same z-score for a certain Pearson correlation, regardless if the number of time points used to estimate the correlation is 50 or 500 (and regardless of the temporal auto correlation in the data). As for task fMRI, where unbiased beta coefficients from the first level analyses are used in the group analysis (and not test statistics like t- or z-scores), the correlation as well as the raw Fisher transformed correlation will be (approximately) unbiased and will therefore lead to nominal results (even for large group differences in motion scrubbing).

The mean FWE rate for CCS is lower compared to CPAC and DPARSF. A possible explanation is that the CCS pipeline includes a despiking step, which will reduce ROI correlations due to spikes (as a spike will be present in most ROIs).

### 5.2. Limitations

The motion scrubbing used in this paper is completely random, which means that low motion time points can be removed while high motion time points are still present in the data. The reason for using this approach is that the ABIDE preprocessed dataset does not contain motion metrics for each time point. In future work we will perform more realistic motion scrubbing, based on different motion metrics, such that high motion time points are the most likely to be removed.

The group analyses are based on using data from both individuals diagnosed with autism spectrum disorder and typical controls. Ideally the group analyses should be performed using only controls, but this will lead to a small number of subjects from each site.

We have here only looked at false positives, but false negatives are also important (Noble et al., 2020). To estimate statistical power using real fMRI data is not a trivial task. For group analyses involving a single group, the brain activity or connectivity of a large cohort of subjects (e.g. 400) can be seen as the ground truth, and power can then be estimated as the number of times a group analysis with a smaller number of subjects (e.g. 20) results in the same significant activity or connectivity (Lohmann et al., 2018). For group analyses involving two groups of subjects (our case), it is necessary to start with an analysis that provides a significant difference between the two groups, or to artificially add a difference and count how many times it is detected (Dansereau et al., 2017). In future work we will investigate how this can be done for resting state fMRI.

In the function nets glm in FSLNets a permutation test is used to perform the group analysis and to obtain corrected p-values. We intend to try the permutation based approach, to see if it reduces the degree of false positives, but without GPU acceleration it is rather time consuming to use for so many group analyses and parameter combinations. We will investigate if BROCCOLI (Eklund et al., 2014) can be used instead of randomise, to speedup the permutations. Furthermore, a bug was detected in nets glm^5^ and we are waiting for this bug to be fixed.

### 5.3. Impact

It is difficult to estimate the impact of our findings, since resting state fMRI entails many different preprocessing choices (e.g. bandpass filtering, global mean regression, nuisance regression) and it is not clear how common different parameters, softwares and pipelines are. For example, it is unknown how common FSLNets (with auto correlation corrected Fisher r-to-z) is compared to REST (Song et al., 2011) and Conn (Whitfield-Gabrieli & Nieto-Castanon, 2012) (both without auto correlation correction). It is also not clear if full correlation is more popular than partial correlation, and how common it is with group analyses where the motion scrubbing substantially differs between two groups. Carp (2012) reviewed methods reporting in the fMRI literature, but did not cover aspects specific to resting state fMRI.

## Acknowledgements

The authors have no conflict of interest to declare. This study was supported by Swedish research council grant 2017-04889 and CENIIT at Linköping University. Funding was also provided by the ITEA3 / VINNOVA funded project “Intelligence based iMprovement of Personalized treatment And Clinical workflow supporT” (IMPACT).

## Footnotes

1 http://preprocessed-connectomes-project.org/abide/

2 http://preprocessed-connectomes-project.org/abide/Pipelines.html

3 https://fsl.fmrib.ox.ac.uk/fsl/fslwiki/FSLNets

4 https://github.com/wanderine/GroupDifferencesMotionScrubbing

5 http://tiny.cc/n0wwjz

